# Genomic Benchmarks: A Collection of Datasets for Genomic Sequence Classification

**DOI:** 10.1101/2022.06.08.495248

**Authors:** Katarina Gresova, Vlastimil Martinek, David Cechak, Petr Simecek, Panagiotis Alexiou

## Abstract

In this paper, we propose a collection of curated and easily accessible sequence classification datasets in the field of genomics. The proposed collection is based on a combination of novel datasets constructed from the mining of publicly available databases and existing datasets obtained from published articles. The main aim of this effort is to create a repository for shared datasets that will make machine learning for genomics more comparable and reproducible while reducing the over-head of researchers that want to enter the field. The collection currently contains eight datasets that focus on regulatory elements (promoters, enhancers, open chromatin region) from three model organisms: human, mouse, and roundworm. A simple convolution neural network is also included in a repository and can be used as a baseline model. Benchmarks and the baseline model are distributed as the Python package ‘genomic-benchmarks’, and the code is available at https://github.com/ML-Bioinfo-CEITEC/genomic_benchmarks.

## Introduction

Recently, deep neural networks have been successfully applied to identify functional elements in the genomes of humans and other organisms, such as promoters Oubounyt et al. (2019), enhancers Le et al. (2021), transcription factor binding sites Quang and Xie (2019), and others. Neural network models have been shown to be capable of predicting histone accessibility Yin et al. (2019), RNA-protein binding Shen et al. (2020), and accurately identify short non-coding RNA loci within the genomic background Georgakilas et al. (2020).

However, deep neural network models are highly dependent on large amounts of high-quality training data Sun et al. (2017). Comparing the quality of various deep learning models can be challenging, as the authors often use different datasets for evaluation, and quality metrics can be heavily influenced by data preprocessing techniques and other technical differences Nawi et al. (2013).

Many computational fields have developed established benchmarks, for example, SQuAD for question answering Rajpurkar et al. (2016), IMDB Sentiment for text classification Maas et al. (2011), and ImageNet for image recognition Deng et al. (2009). Benchmarks are crucial in driving innovation. The annual competition for object identification Russakovsky et al. (2015) catalyzed the boom in AI, leading in just seven years to models that exceed human capabilities.

In biology, a great challenge over the past 50 years has been *the protein folding problem*. To compare different protein folding algorithms, the community introduced the Critical Assessment of protein Structure Prediction (CASP) Moult et al. (1995) challenge benchmark that provides research groups with the opportunity to objectively test their methods. In 2021, AlphaFold Jumper et al. (2021) won this competition producing predicted structures within the error tolerance of experimental methods. This carefully curated benchmark led to the solution of the most prominent bioinformatic challenge of the past 50 years.

In Genomics, we have similar challenges like annotation of genomes and identification and classification of functional elements, but currently we lack benchmarks similar to CASP. Practically, machine learning tasks in Genomics commonly involve the classification of genomic sequences into several categories and/or contrasting them to a genomic background (a negative set). For example, a well-studied question in Genomics is the prediction of enhancer loci on a genome. For this question, the benchmark situation is highly fragmented. As an example, Liu et al. (2016) proposed a benchmark dataset based on the chromatin state from multiple cell lines. Both enhancer and non-enhancer sequences were retrieved from experimental chromatin information. The CD-HIT software Li and Godzik (2006) was used to filter similar sequences, and the benchmark dataset was made available as a pdf file. However, information stored in a pdf file is suitable for human communication, but computers cannot easily extract data from these files. Despite not being easily machine readable, it was used by many subsequent publications (Liu et al. (2018), Le et al. (2019), Tahir et al. (2017), Jia and He (2016), He and Jia (2017), Nguyen et al. (2019), Khanal et al. (2020), Le et al. (2021), Zhang et al. (2021), Inayat et al. (2021), Mu et al. (2021) or Yang et al. (2021)) as a gold standard for enhancer prediction, highlighting the need for benchmark datasets in this field. Other common sources of enhancer data are the VISTA Enhancer Browser Visel et al. (2007), the FANTOM5 Andersson et al. (2014), the ENCODE project ENCODE Project Consortium et al. (2012), and the Roadmap Epigenomics Project Kundaje et al. (2015) which provide a wealth of positive samples but no negatives. A researcher would need to implement their own method of negative selection, thus introducing individual selection biases to the samples.

Another highly studied question in Genomics is the prediction of promoters. Benchmark situation in this field has its own problems. For example, Lin and Li (2011) extracted positive samples from EPD Schmid et al. (2006) and the non-promoter sequences were randomly extracted from coding regions and non-coding regions, and used as two negative sets. This method for creating a negative set is not an established one. Other authors used only coding sequences or only non-coding sequences as a negative set Gordon et al. (2003) or combined coding and non-coding sequences as a one negative set Ohler (2006), Yang et al. (2008), Rani et al. (2007). Even Lin and Li (2011) are already pointing to the problem of missing benchmarks and reproducibility, saying that it is difficult to compare their results with other published results due to differences in data and experimental protocol. Several years later, Lai et al. (2019) created their own dataset and reported similar problems. They were unable to compare the results with other published tools because the datasets were derived from different sources, used different proprocessing procedures, or were not made available at all.

In this paper, we propose a collection of benchmark datasets for the classification of genomic sequences, focusing on ease of use for machine learning purposes. The datasets are distributed as a Python package ‘genomic-benchmarks’ that is available on GitHub^1^ and distributed through The Python Package Index (PyPI)^2^. The package provides an interface that allows the user to easily work with the benchmarks using Python. Included are utilities for data processing, cleaning procedures, and summary reporting. Additionally, it contains functions that make training a neural network classifier easier, such as PyTorch Paszke et al. (2019) and TensorFlow Abadi et al. (2016) data loaders and notebooks containing basic deep learning architectures that can be used as templates for prototyping new methods. Importantly, every dataset presented here comes with an associated notebook that fully reproduces the dataset generation process, to ensure transparency and reproducibility of benchmark generation in the future.

## Results and Methods

### Overview of Datasets

The currently selected datasets are divided into three categories. There is a group of datasets focused on human regulatory functional elements, either produced from mining the Ensembl database, or from published datasets used in multiple articles. For promoters, we have imported human non-TATA promoters Umarov and Solovyev (2017). For enhancers, we used human enhancers from Cohn et al. (2018) paper and Ensembl human enhancers from the FANTOM5 Project Andersson et al. (2014). We have also included open chromatin regions and multiclass dataset composed of three regulatory elements (enhancers, promoters, and open chromatin regions), both constructed from the Ensembl regulatory build Zerbino et al. (2015). The second category consists of ‘demo’ datasets that were computationally generated for this project, and focus on classification of genomic sequences between different species or types of transcripts (protein coding vs non-coding). Finally, the third category ‘dummy’ has a single small dataset which can be used for quick prototyping of methods due to its small size. From the point of view of model organism, our datasets include primarily human data, but also mouse (*Mus musculus*), and roundworm (*Caenorhabditis elegans*). An overview of available datasets is given in Table 1 and simple code for listing all currently available datasets in Figure 1.

**Table 1.** Description of datasets in genomic benchmark package. Several pieces of information are provided about each dataset: a) *Name* is unique identification of dataset in genomic benchmark package b) *# of sequences* is combined count of all sequences from all classes c) *# of classes* is count of all classes in a dataset d) *Class ratio* is a ratio between number of sequences in a biggest class and number of sequences in a smallest class e) *Median length* is computed for all sequences from all classes in a dataset f) *Standard deviation* is also computed for all sequences from all classes in a dataset.

| Name | # of sequences | # of classes | Class ratio | Median length | Standard deviation |
| --- | --- | --- | --- | --- | --- |
| dummy_mouse_enhancers_ensembl | 1210 | 2 | 1.0 | 2381 | 984.4 |
| demo_coding_vs_intergenomic_seqs | 100000 | 2 | 1.0 | 200 | 0.0 |
| demo_human_or_worm | 100000 | 2 | 1.0 | 200 | 0.0 |
| human_enhancers_cohn | 27791 | 2 | 1.0 | 500 | 0.0 |
| human_enhancers_ensembl | 154842 | 2 | 1.0 | 269 | 122.6 |
| human_ensembl_regulatory | 289061 | 3 | 1.2 | 401 | 184.3 |
| human_nontata_promoters | 36131 | 2 | 1.2 | 251 | 0.0 |
| human_ocr_ensembl | 174756 | 2 | 1.0 | 315 | 108.1 |

**Fig. 1.**
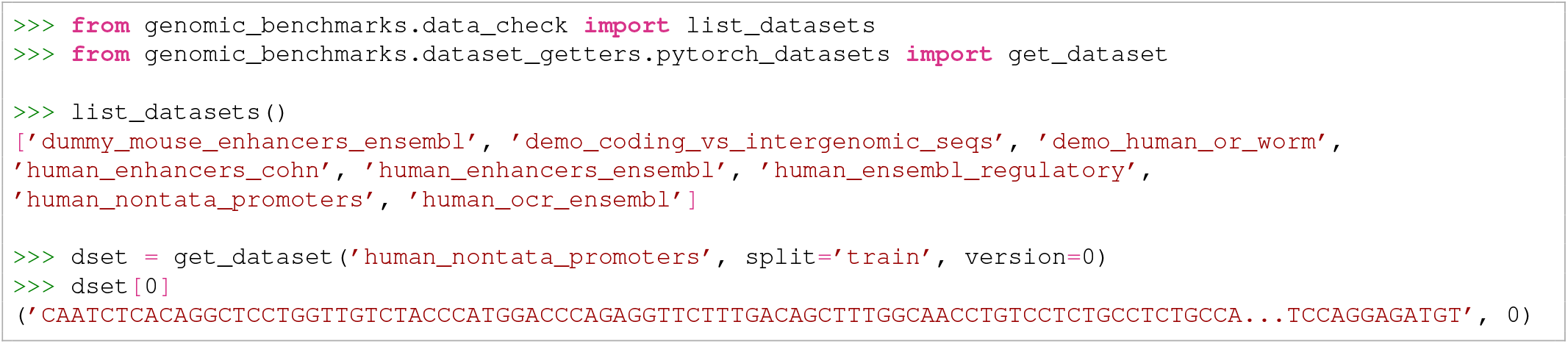
Listing all available datasets and getting PyTorch Dataset for one of them.

The *Human enhancers Cohn* dataset was adapted from Cohn et al. (2018). Enhancers are genomic regulatory functional elements that can be bound by specific DNA binding proteins so as to regulate the transcription of a particular gene. Unlike promoters, enhancers do not need to be in a close proximity to the affected gene, and may be up to several million bases away, making their detection a difficult task.

The *Human enhancers Ensembl* dataset eas constructed from Human enhancers from The FANTOM5 project Andersson et al. (2014) accessed through the Ensembl database Howe et al. (2021). Negative sequences have been randomly generated from the Human genome GRCh38 to match the lengths of positive sequences and not overlap them.

The *Human non-TATA promoters* dataset was adapted from Umarov and Solovyev (2017). These sequences are of length 251bp: from -200 to +50bp around transcription start site (TSS). To create non-promoters sequences of length 251bp, the authors of the original paper used random fragments of human genes located after first exons.

The *Human ocr Ensembl* dataset was constructed from the Ensembl database Howe et al. (2021). Positive sequences are Human Open Chromatin Regions (OCRs) from The Ensembl Regulatory Build Zerbino et al. (2015). Open chromatin regions are regions of the genome that can be preferentially accessed by DNA regulatory elements because of their open chromatin structure. In the Ensembl Regulatory Build, this label is assigned to open chromatin regions, which were experimentally observed through DNase-seq, but covered by none of the other annotations (enhancer, promoter, gene, TSS, CTCF, etc.). Negative sequences were generated from the Human genome GRCh38 to match the lengths of positive sequences and not overlap them.

The *Human regulatory Ensembl* dataset was constructed from Ensembl database Howe et al. (2021). This dataset has three classes: enhancer, promoter and open chromatin region from The Ensembl Regulatory Build Zerbino et al. (2015). Open chromatin region sequences are the same as the positive sequences in the Human ocr Ensembl dataset.

### Reproducibility

The pre-processing and data cleaning process we followed is fully reproducible. We provide a Jupyter notebook that can be used to recreate each given dataset, and can be found in the `docs` folder of the GitHub repository^3^. All dependencies are provided, and a fixed random seed is set so that the notebook will always produce the same data splits. Each dataset is divided into training and testing subsets. For some datasets, which contain only positive samples, we had to generate appropriate negative samples (dummy mouse enhancers Ensembl, human enhancers Ensembl and human open chromatin region Ensembl dataset). These negatives were randomly selected from the same genome as the positive samples. The length distribution of positive samples was matched when selecting the negatives, and regions overlapping with the positive samples were excluded from the selection.

### Data format

All samples were stored as genomic coordinates, and datasets originally provided as sequences (human enhancers Cohn, human nonTATA promoters) were mapped to the reference using the ‘seq2loc‘ tool included in the package. Data were stored as compressed (gzipped) CSV tables of genomic coordinates, containing all information typically found in a BED format table. Column names are *id, region, start, end*, and *strand*. Each dataset has *train* and *test* sub-folders and a separate table for each class. Furthermore, each dataset contains a YAML information file with metadate such as its version, the names of included classes, and links to sequence files of the reference genome. The stored coordinates and linked sequence files were used to produce the final datasets, ensuring the reproducibility of our method. For more information, visit the `datasets` folder of the GitHub repository^4^. To speed up this conversion from a list of genomic coordinates to a locally stored folder of nucleotide sequences, we provide a cloud based cache of the full sequence datasets which can be used simply by setting the `use_cloud_cache=True option`.

### Easy data access tools

We provide ready-to-use data loaders for the two most commonly used deep learning frameworks, TensorFlow and PyTorch. These data loaders allow the user to load any of the provided datasets using a single line of code (for example Figure 1). This feature is important for reproducibility and for adoption of the package, particularly by people with limited knowledge of genomics. Moreover, an example of usage for a simple convolutional neural network (adapted from Klimentova et al. (2020)) is provided in the `notebooks` folder of the GitHub repository^5^. The neural network consists of three convolutional layers with 16, 8, and 4 filters, with a kernel size of 8. The output of each convolutional layer goes through the batch normalization layer and the max-pooling layer. The output of the last set of layers is flattened and goes through two dense layers. The last layer is designed to predict probabilities that the input sample belongs to any of the given classes. The architecture of the model is shown in Figure 2. To get a baseline estimate for researchers using these benchmarks, we fit the convolutional neural network model described above to each dataset included in our collection. Training notebooks are provided in an `xperiments` folder of the GitHub repository^6^. The models were trained for 10 epochs with batch size 64. The accuracy and F1 score for PyTorch and Tensorflow CNN models on all genomic benchmark datasets are shown in Table 2.

**Fig. 2.**
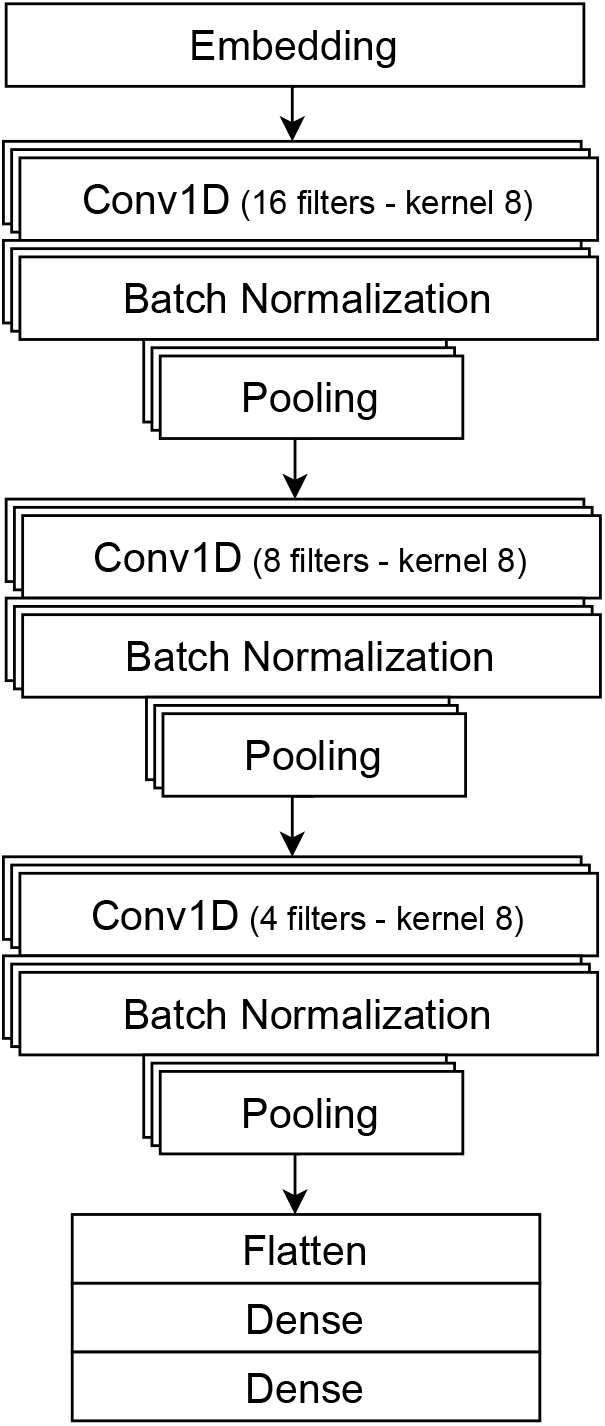
CNN architecture. The neural network consists of three convolutional layers with16, 8, and 4 filters, with a kernel size of 8. The output of each convolutional layer goes through the batch normalization layer and the max-pooling layer. The output is then flattened and passes through two dense layers. The last layer is designed to predict the probabilities that the input sample belongs to any of the given classes.

**Table 2.** Performance of baseline models on benchmark datasets.

| Dataset | Pytorch |  | Tensorflow |  |
| --- | --- | --- | --- | --- |
|  | Accuracy | F1 score | Accuracy | F1 score |
| dummy_mouse_enhancers_ensembl | 69.0 | 70.4 | 50.0 | 66.9 |
| demo_coding_vs_intergenomic_seqs | 87.6 | 86.8 | 89.6 | 89.4 |
| demo_human_or_worm | 93.0 | 92.8 | 94.2 | 93.2 |
| human_enhancers_cohn | 69.5 | 67.1 | 68.9 | 71.3 |
| human_enhancers_ensembl | 68.9 | 56.5 | 81.1 | 74.6 |
| human_ensembl_regulatory | 93.3 | 93.3 | 79.3 | 79.3 |
| human_nontata_promoters | 84.6 | 83.7 | 86.5 | 84.4 |
| human_ocr_ensembl | 68.0 | 66.1 | 68.8 | 72.0 |

## Discussion

Machine learning, and especially deep learning, techniques have recently started revolutionizing the field of genomics. Deep learning methods are highly dependent on large amounts of high-quality data to train and benchmark data are needed to accurately compare performance of different models. Here, we propose a collection of Genomic Benchmarks, produced with the aim of being easily accessible and reproducible. Our intention is to lower the difficulty of entry into the machine learning for Genomics field for researchers that may not have extensive knowledge of Genomics but want to apply their knowledge of machine learning in this field. Such an approach worked well for the field of protein folding, where benchmark-based competitions helped revolutionize the field.

The eight genomics datasets that have been currently added are a first step towards the direction of a large repository of Genomic Benchmarks. Beyond making access to these datasets easy for users, we have ensured that adding more datasets in a reproducible way is an easy task for further development of the repository. We encourage users to propose datasets or subfields of interest that would be useful in future releases. We have provided guidelines and tools to unify access to any genomic data and we will happily host submitted genomic datasets of sufficient quality and interest.

We are aware of the limitations of the current repository. While we strive to include diverse data, still most of our benchmark datasets are balanced, or close to balanced, having similar length of sequences and a limited number of classes. Our main datasets all come from the human genome, and all deal with regulatory features. In the future, we would like to increase the diversity of our datasets to be able to diagnose the model’s sensitivity to those factors. Many machine learning tasks in Genomics consist of binary classification of a class of Genomic functional elements against a background. However, it can be beneficial to start expanding the field into multi-class classification problems, especially for functional elements that have similar characteristics to each other against the background. We will expand our benchmark collection to include more imbalanced datasets, and more multi-class datasets.

In this manuscript, we have implemented a simple convolutional neural network as a baseline model trained and evaluated on all of our datasets. Improvement on this baseline will be certainly achieved by using different architectures and training schemes. We have an open call for users that out-perform the baseline to submit their solution via our Github repository, and be added to a ‘Leaderboard’ of methods for each dataset. We hope that this will create a healthy competition on this set of reproducible datasets, and promote machine learning research in Genomics.

## ACKNOWLEDGEMENTS

We are thankful to Google Cloud for providing P. Simecek and V. Martinek free research credits. Some computational resources were supplied by the project “e-Infrastruktura CZ” (e-INFRA CZ LM2018140) supported by the Ministry of Education, Youth and Sports of the Czech Republic.

## AUTHOR CONTRIBUTIONS

KG did current state of the field research. KG and PS created and collected datasets. VM implemented data loaders. DC, PS and KG implemented baseline models. KG, PS and PA prepared the manuscript. All authors read and approved the final manuscript.

## Footnotes

1 https://github.com/ML-Bioinfo-CEITEC/genomic_benchmarks

2 https://pypi.org/project/genomic-benchmarks/

3 https://github.com/ML-Bioinfo-CEITEC/genomic_benchmarks/tree/main/docs

4 https://github.com/ML-Bioinfo-CEITEC/genomic_benchmarks/tree/main/datasets

5 https://github.com/ML-Bioinfo-CEITEC/genomic_benchmarks/tree/main/notebooks

6 https://github.com/ML-Bioinfo-CEITEC/genomic_benchmarks/tree/main/experiments

